# Mechanochemical signal transduction in synthetic cells

**DOI:** 10.1101/2022.04.26.489423

**Authors:** Kevin Jahnke, Maja Illig, Marlene Scheffold, Mai P. Tran, Ulrike Mersdorf, Kerstin Göpfrich

## Abstract

Mechanotransduction determines the adaptive response of natural cells via transmembrane proteins^1^. The incorporation of membrane-spanning structures to guide cellular function and to enable transmembrane signalling is therefore a critical aim for bottom-up synthetic biology^2,3,4^. Here, we design membrane-spanning DNA origami signalling units (DOSUs) and mechanically couple them to DNA cytoskeletons^5^ encapsulated within giant unilamellar vesicles (GUVs). We verify the assembly and incorporation of the DOSUs into the GUV membranes and achieve their clustering upon external stimulation. The transmembrane-spanning DOSUs act as a pore to allow for the transport of single-stranded DNA into the GUVs. We employ this to externally trigger the reconfiguration of DNA cytoskeletons within GUVs using strand displacement reactions. In addition to chemical signalling, we achieve the mechanical coupling of the externally added DOSUs and the internal DNA cytoskeletons. We induce clustering of the DOSUs, which triggers a symmetry break in the organization of the DNA cytoskeleton which is mechanically coupled to the DOSU.Our work thus provides a mechanical and chemical transmembrane signaling module towards the assembly of stimuli-responsive and adaptive synthetic cells.

## Main

DNA nanotechnology emerged as a powerful tool for bottom-up synthetic biology^6^. In particular, versatile DNA nanostructures have been engineered to function as transmembrane pores which allow for the transport of ions^7,8,9^, fluorophores^10,11^, macromolecules^12,13^, DNA^14^, proteins^15^ or lipids^16^. However, apart from molecular substrate exchange, a mechanochemical connection to intracellular downstream functions within synthetic cellular compartments is missing. Concomitantly, DNA nanotubes as structural and functional mimics of cytoskeletons have been reconstituted within cell-sized water-in-oil droplets^17,18^ and recently also in GUVs^5^. Here, we link filament disassembly and cytoskeletal remodeling within GUVs to transmembrane properties of DNA origami and chemical signalling from the outside of GUVs reminiscent of focal adhesions within natural cells^19^. Thereby, we advance the current engineering of synthetic cells with synthetic cytoskeletons and transmembrane DNA signalling units by coupling their mechanochemical properties across the compartment barrier.

First, we set out to design a transmembrane-spanning DNA origami signalling unit (DOSU) with the following functional features: (i) The DOSUs should have the ability to act as nanopores for the translocation of small molecules. (ii) It should be possible to cluster the DOSUs to mimic receptor clustering as a transmembrane signalling mechanism. (iii) They should enable chemical signalling across the GUV membrane. (iv) The transmembrane-spanning DOSUs should be mechanically coupled to a cytoskeleton within the GUV to mimic focal adhesions. Taken together, we thus aim for mechanochemical signal transduction across the GUV membrane. Fig. 1**a** illustrates the basic design of our transmembrane DOSU. We adapt a DNA origami design which was initially used for nucleating algorithmic self-assembly^20^ and later as a binding site for DNA nanotubes, building superstructures and junctions^21,22,23,24,25,26^. We modified this DNA origami with 12 overhangs that can each bind to a cholesterol-tagged DNA (Supplementary Fig. 1**a**). This allows the attachment and insertion of the DOSU into lipid membranes. Additionally, the design contains 100 fluorophore modifications to aid the visualization of individual DOSUs with confocal microscopy (Supplementary Fig. 1**c**). First, we verify the successful assembly of the DOSU with confocal microscopy (Supplementary Fig. 2) and agarose gel electrophoresis (Supplementary Fig. 3). They remain stable for more than three days after purification and even in the absence of magnesium ions (Supplementary Fig. 4). Next, we test the fist functional feature of our DOSU, namely its ability to act as a transmembrane pore (Feature (i)). By incubating DOSUs with cholesterol-tagged DNA and subsequently mixing them with SUVs with a diameter of 100 nm, we observe the binding of the DOSUs to the SUV membrane via cryo electron microscopy (cryo-EM, Fig. 1**b**, Supplementary Fig. 5). If the DOSU is penetrating the membrane as designed, it should be bound at a perpendicular angle to the SUV surface. By analyzing the angle of bound DOSUs to the SUV membrane, we find that 87 %indeed attach to the SUV membrane in the desired orientation (75-105°, Fig. 1**c**, Supplementary Fig. 6). To further prove their insertion into the lipid membrane, we conduct dye influx assays. To enable vizualization with confocal microscopy, we bind the DOSUs to giant unilamellar vesicles (GUVs) (Fig. 1**d**) containing a low fraction of fluorescently labelled lipids (0.5% Atto488-PE) and add a water-soluble membrane impermeable Alexa647-NHS ester fluorescent analyte from the outside. With the DNA origami concentration measured with UV-vis spectroscopy, we approximate the number of DOSUs on the GUV membrane to be 10700 ±1200, which corresponds to a density of (16 ±1) DOSUs/μm^2^(Supplementary Note, Supplementary Fig. 7). Following the addition of the fluorescent analyte, we observe the dye influx over time in presence and absence of the DOSUs (Fig. 1**e**). To quantify the amount of dye influx, we analyze the intensity ratio inside and outside of the GUVs over time. The amount of fully-permabilized GUVs increases over time for GUVs containing the DOSUs (Fig. 1**f**). On the other hand, no dye influx can be observed in their absence or in absence of cholesterol tags (Supplementary Fig. 8). After 330 min, the median value of the ratio I_in_ /I_out_ is 0.99 in presence of DOSUs and 0.08 in the absence, respectively. This confirms the successful engineering of transmembrane DOSUs that can serve as pore for small molecules like fluorophores (Feature (i)). Importantly, the median value of the intensity ratio, and thereby fraction of fully-permeabilized GUVs over unpermeabilized GUVs, can be tuned by changing the concentration and density of DOSUs on the membrane (Supplementary Fig. 9). Note that we can exclude GUV permeabilization due to osmotic stress, as we add the same amount of iso-osmotic solutions for all conditions and beyond we can assume that the DOSU-mediated membrane permeation is due to a non-transient insertion (Supplementary Fig. 10). In addition to the dye influx assay, we also observe single GUVs over time and perform dye efflux assays. Fluorophore transport happens within minutes after signalling unit insertion (Supplementary Fig. 11, Supplementary Video 1).

**Figure 1:**
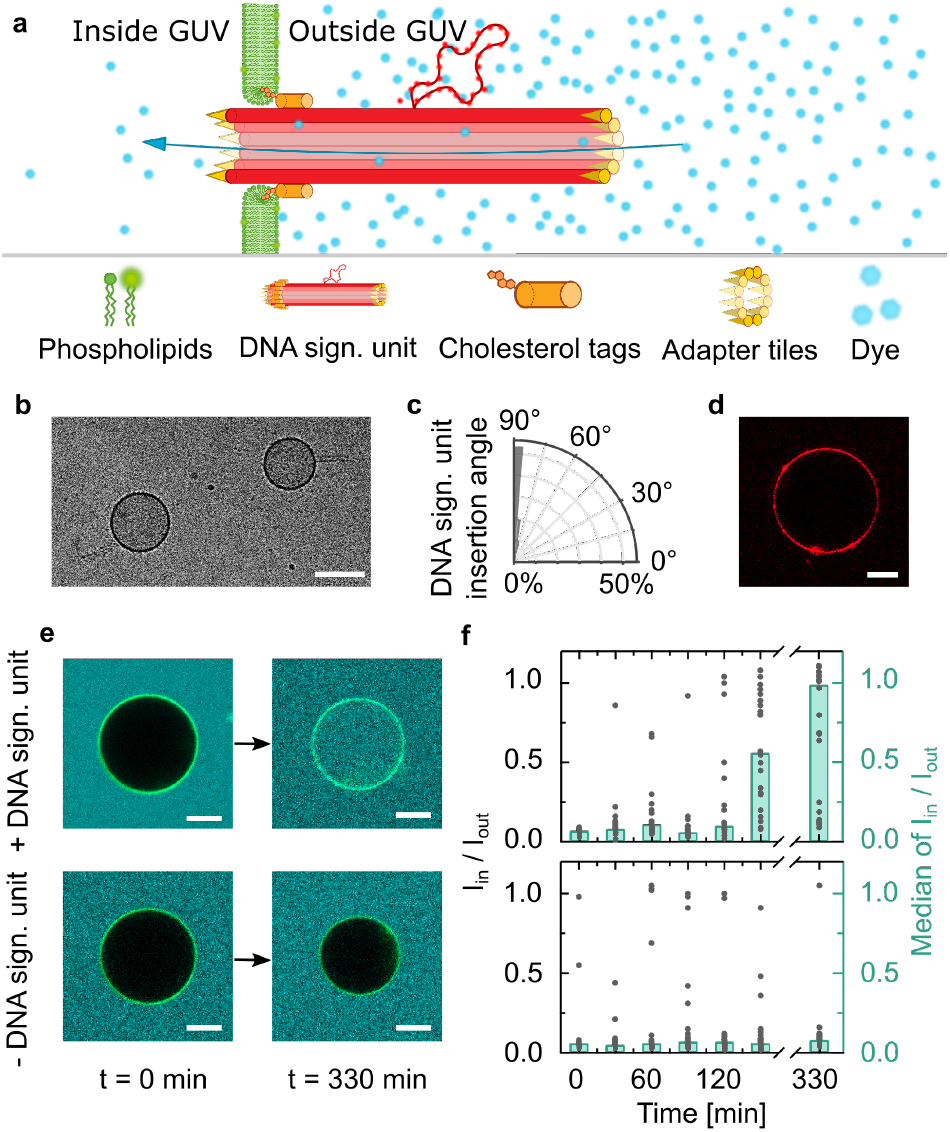
Cholesterol-tagged DNA origami signalling units (DOSUs) insert into phospholipid bilayers. **a** Schematic representation of the cylindrical DNA origami signalling unit design inserted into the GUV membrane via 12 cholesterol tags. The DOSU creates a passage for fluorophores to be transported into the GUV lumen. **b** Representative cryo electron micrograph of small unilamellar vesicles (SUVs) with transmembrane DOSUs. Scale bar: 100 nm. **c** Insertion angle of the DOSU in SUVs extracted from cryo-EM micrographs (*n* = 287). **d** Confocal image of DOSUs (red, Atto647-labeled, *λ*_ex_ = 640 nm) inserted into a GUV membrane. Scale bar: 10 μm. **e** Confocal images of GUVs (green, Atto488-labeled, *λ*_ex_ = 488 nm) immersed in a solution containing a fluorescent analyte (cyan, Atto647, *λ*_ex_ = 640 nm). The first row shows GUVs with DOSUs and the second row without DOSUs at t =0 min and t =330 min, respectively. Scale bars: 10 μm. **f** Ratio of inner and outer intensity I_in_/I_out_*of* n ≥ 20 individual GUVs over time and histogram of the median value of *I*_in_/*I*_out_ with (top, n ≥ 20) and without (bottom, n ≥ 30) DOSUs.

A remarkable feature of transmembrane proteins is the ability to cluster in response to a molecular trigger to enable a certain function, e.g. the formation of focal adhesions^19^. To engineer the clustering of our DOSUs (Feature (ii)), we modified 6 DNA oligos with biotin around the DNA origami circumference (Fig. 2**a**, Supplementary Fig. 1**b**). Streptavidin serves as a stimulus to induce DOSU clustering in bulk (Fig. 2**b**, Supplementary Fig. 12) and on the GUV membrane (Fig. 2**c**) as we confirmed with confocal microscopy. The degree of clustering is increasing for increasing streptavidin concentrations within the range from 0 μM to 4 μM for a given DOSU concentration (100 ng/μL of DNA). Interestingly, dye influx experiments indicate that the membrane permeabilization is enhanced for clustered DOSUs compared to unclustered DOSUs (Fig. 2**d**). The median value of *I*_in_/*I*_out_ 30 min and 60 min after addition of streptavidin is 0.24, 0.54 for clustered and 0.13, 0.19 for unclustered DOSUs, respectively.

**Figure 2:**
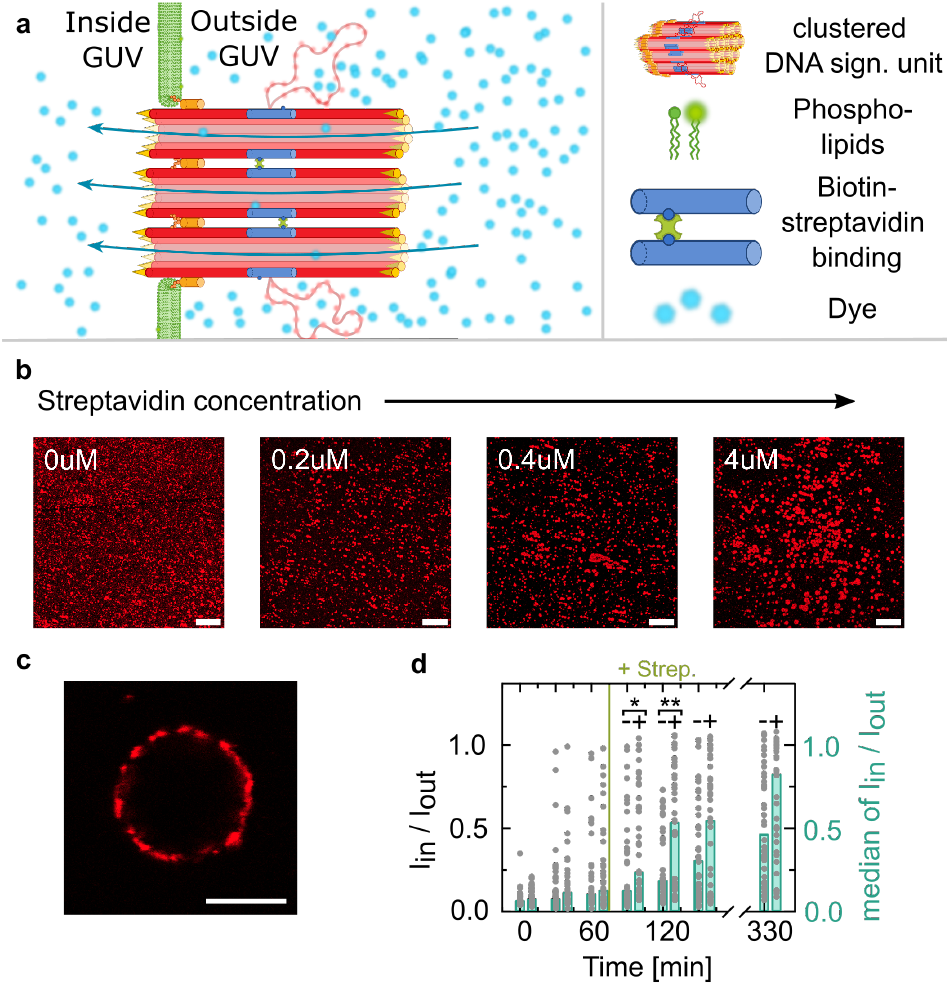
Stimuli-induced DOSU clustering. **a** Schematic representation of biotinylated DOSUs inserted into a GUV membrane and clustered upon addition of streptavidin. **b** Confocal images of clustered DOSUs (red, Atto647-labeled, *λ_ex_* = 640 nm) with increasing streptavidin concentration, forming visible clusters from 0.2 to 4 μM streptavidin. Scale bars: 10 μm. **c** Confocal image of a GUV with clustered DOSUs (red, Atto647-labeled, *λ_ex_* = 640 nm) inserted into the membrane. Scale bar: 10 μm. **d** Ratio of inner over outer intensity *I*_in_/*I*_out_ of individual GUVs (n ≥ 40, two technical repeats) over time and histogram of the median value of *I*_in_/*I*_out_ for GUVs without strepatvidin and GUVs 60 min after the addition of streptavidin. The dye influx is significantly increased after addition of streptavidin indicating better insertion and transport rates for clustered DOSUs. A Mann-Whitney test was performed, the p-values are 0.019 and 0.007 for *t* = 90 min and *t* = 120 min, respectively.

A second major function of natural pores is the transport of essential molecules across the membrane for chemical transmembrane signalling. To employ the engineered DOSUs for chemical signalling (Feature (iii)), we interfaced our DNA pores with DNA-based mimics of cytoskeletons. We used DNA double-crossover tiles^27^ which self-assemble into hollow DNA filaments and encapsulated them into GUVs^5^. The DNA filaments were modified with single-stranded overhangs so that they can be disassembled by toehold-mediated strand displacement^18^. The presence of an invader DNA strand thereby leads to filament disassembly by binding the overhang of a single DNA tile (Supplementary Video 2). By transporting the invader strand across the GUV membrane via the clustered DOSUs, we thus aim to create a chemical signalling pathway, whereby an external stimulus leads to filament disassembly in the interior of the GUV (Fig. 3**a**). First, we verify that the DOSUs can transport fluorescently-labeled single-stranded DNA (ssDNA) across the GUV membrane. Over the course of 330 min, we observe that the median value of *I*_in_/*I*_out_ reaches 0.75 (Fig. 3**b** and 3**c**). The smaller amount of fully-permeabilized GUVs at this time point is in agreement with the bigger size and slower diffusion of ssDNA in comparison to the fluorescent analyte. Notably, ssDNA influx was also achieved with the unclustered signalling unit (Supplementary Fig. 13), which is consistent with the outer diameter of the pore (12 nm). We can thus translocate the invader strand into the GUV lumen, which is filled with DNA filaments. In presence of DOSUs, we observe the filament disassembly inside GUVs over time (Fig. 3**d**). As a measure for the degree of filament disassembly, we quantify the porosity of the GUV lumen containing DNA filaments from confocal images (see Methods and Supplementary Fig. 14). After 5 h of incubation with streptavidin, the normalized porosity decreases from 1 to 0.53 ±0.18 in presence of the DOSUs (Fig. 3**e**). As we know from the influx experiments with fluorescently labelled DNA (Supplementary Fig. 13), the GUVs are fully permeabilized at this time point. Moreover our results indicate that filament disassembly for clustered DOSUs is enhanced compared to unclustered DOSUs (Supplementary Fig. 15). This shows the successful filament disassembly using chemical signalling across the GUV membrane.

**Figure 3:**
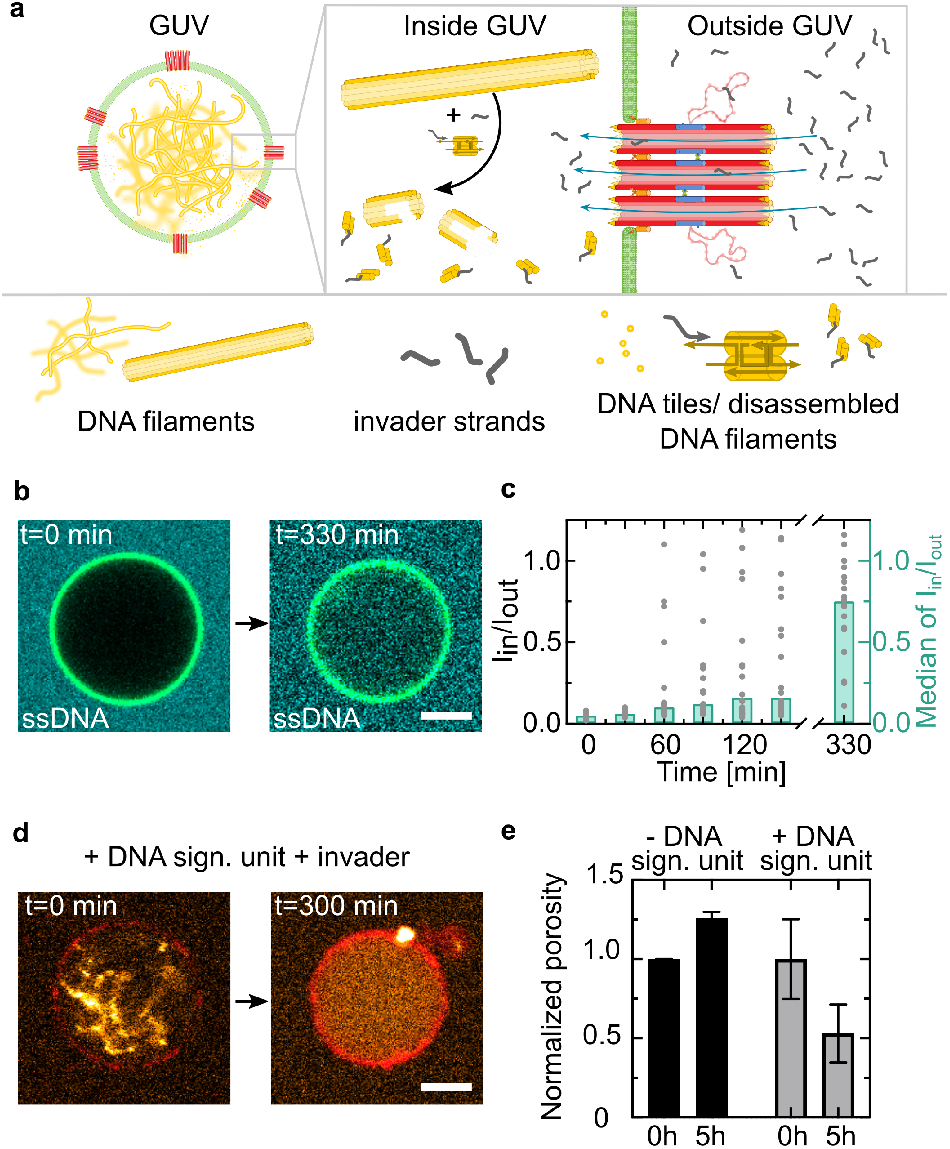
Chemical signal transduction across the GUV membrane. **a** Schematic representation of the influx of single-stranded invader DNA strands through clustered DOSUs to induce disassembly of reconstituted DNA cytoskeletons. **b** Confocal images of GUVs (green, Atto488-labeled, *λ_ex_* =488 nm) with clustered DOSUs immersed in an ssDNA-containing solution (cyan, Atto647-labeled, *λ_ex_* =640 nm) at *t* = 0 min and *t* = 330 min. Scale bars: 10 μm. **c** Ratio of inner over outer intensity *I*_in_/*I*_out_ of individual GUVs over time and histogram of the median value of *I*_in_/*I*_out_ (*n* = 20). **d** Confocal images of GUVs with clustered DOSUs (red, Atto647, *λ_ex_* =640 nm) and encapsulated DNA cytoskeletons (orange, Cy3-labeled, *λ_ex_* =561 nm) immersed in a DNA invader-containing (5 μM) solution. **e** Normalized porosity of DNA cytoskeletons within GUVs after 0 h and 5 h in presence and absence of DOSUs (2 technical repeats, total n ≥ 28, mean ±SD).

Ultimately, we set out to also mechanically couple the externally induced DOSU clustering to filament remodeling inside the GUVs (Feature (iv)). For this purpose, we make use of the cylindrical shape of the DOSU which was designed specifically to allow for the binding to DNA filaments (Fig. 4**a**)^25^. We verify the successful functionalization of the DOSU with confocal microscopy (Fig. 4**b**) and find that after 2 h of incubation, already 39 %of filaments bind to the DOSU for either pre-annealed DNA filaments and also un-annealed DNA tiles. Successful binding increases to 85 %after 22 h (Fig. 4**c**). We can thus link DNA filaments reconstituted on the inside of GUVs to DOSUs added externally due to the transmembrane-spanning nature of the DOSU. After 5 h of incubation, we observe DNA filament remodeling, inducing a symmetry break in the distribution of the filaments in presence of streptavidin (Fig. 4**d**). This showcases the mechanical coupling across the GUV membrane upon the addition of an external chemical stimulus (Supplementary Video 3). To analyze the amount of filament remodeling, we quantify the normalized center-of-mass displacement *r*/*r*_0_ of the filament fluorescence inside GUVs (see Methods, Fig. 4**e**). In presence of streptavidin, the center-of-mass displacement *r*/*r*_0_ normalized to the radius of the GUV is 0.63 ±0.21, whereas it is only 0.20 ±0.10, in the absence of streptavidin. We thereby show how two separate modules for synthetic cells, namely cytoskeletons and transmembrane signalling units, can be combined to realize simplified focal adhesion mimics.

**Figure 4:**
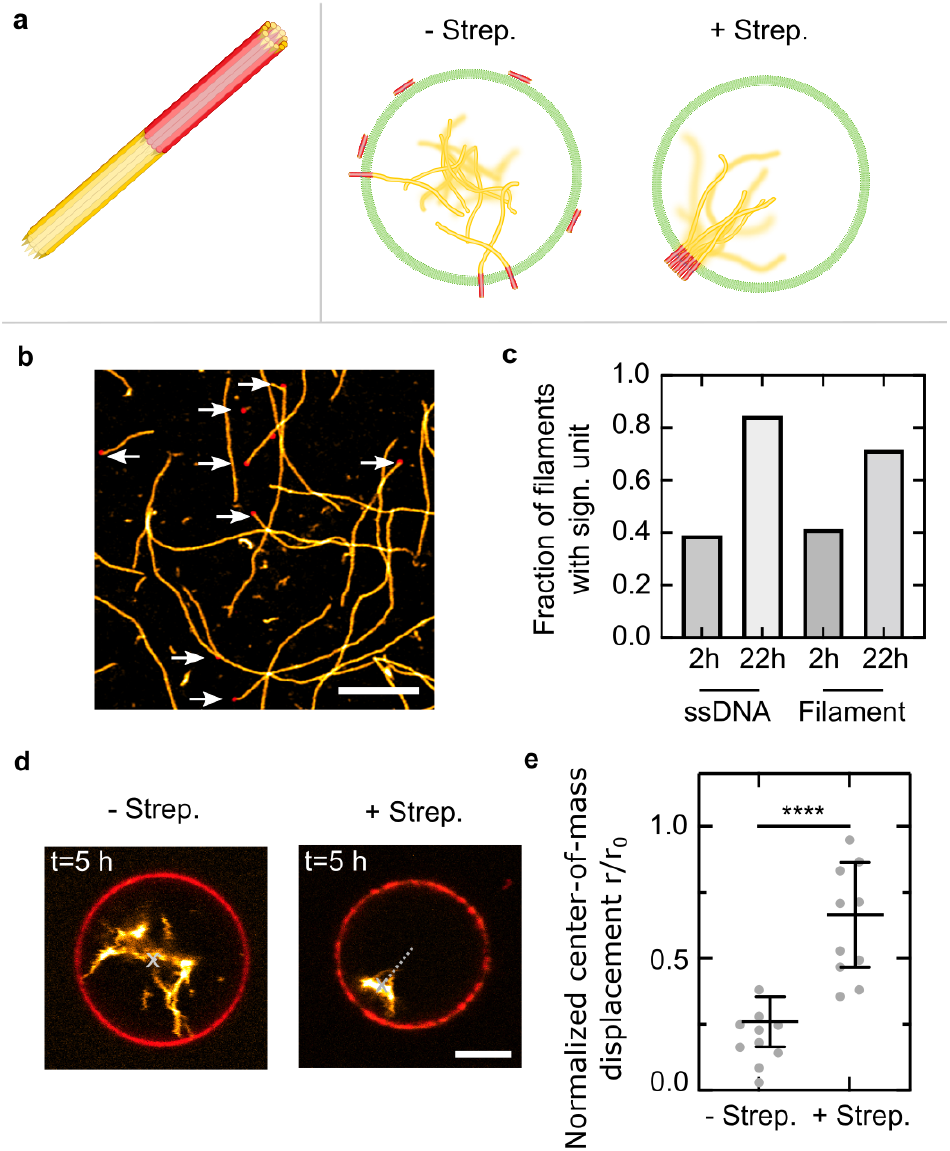
Chemo-mechanical signalling cascade across the GUV membrane induces symmetry breaking of DNA cytoskeletons. **a** Schematic representation of the DOSU attaching to DNA filaments (left). DOSU clustering induces the reorganization of DNA cytoskeletons inside GUVs (right). **b** Confocal image of DOSUs (red, Atto647-labeled, *λ_ex_* = 640 nm) serving as attachment site for DNA filaments (orange, Cy3-labeled, *λ_ex_* = 561 nm). The binding sites are indicated with white arrows. Scale bar: 5 μm. **c** Fraction of filaments bound to DOSUs after 2 h and 22 h for preannealed DNA filaments (‘‘Filament”, n = 628 and n = 162, respectively) and individual tiles that were incubated with the DOSU without annealing (“ssDNA”, n = 186 and n = 196, respectively). **d** Confocal image of DNA cytoskeletons (orange, Cy3-labeled, *λ_ex_* = 561 nm) within GUVs with DOSUs (red, Atto647-labeled, *λ_ex_* = 640 nm) in presence (4 μM) and absence of streptavidin after 5 h of incubation. The center-of-mass of the DNA cytoskeletons within the GUV is indicated. Scale bar: 5 μm. **e** Center-of-mass displacement *r*/*r*_0_ of DNA cytoskeletons within GUVs (normalized to the GUV radius) in presence (4 μM) and absence of streptavidin (*n* = 10, mean ±SD, *p* < 0.0001).

In conclusion, every natural cell interacts with its environment or neighbouring cells using chemical and mechanical signalling pathways across the compartment barrier. This is mainly achieved with transmembrane proteins that can translate external cues into intracellular instructions^28^. Due to its inherent complexity, the reconstitution of natural signalling pathways into synthetic cells is challenging. Moreover, it is especially exciting to build a completely synthetic and rationally engineerable transmembrane mimic that can guide downstream signal transduction in synthetic cells. Here, we designed mechanochemical signal transduction pathways using DNA nanotechnology. We engineer transmembrane DNA origami signalling units that self-assemble into the lipid bilayer of GUVs and can transport small molecules like fluorophores and larger macromolecules, like ssDNA into the GUV lumen. The clustering of our artificial receptors can be triggered by addition of streptavidin as an externally added protein-based stimulus. By reconstituting DNA cytoskeletons into GUVs we implement chemical signalling that induces the filament disassembly upon transport of an invader strand. By binding the DNA filaments to the DOSUs, we also show the mechanical remodeling of the DNA cytoskeleton inside the GUV upon a chemical stimulus from the outside, which clusters the DOSUs. We thereby rationally coupled two completely separate modules for synthetic biology, which was possible by making use of DNA as engineerable molecular building block. In future work, it will be exciting to direct the cell-to-cell communication in between two or more GUVs. Additionally, the engineering of synthetic mechanotransduction pathways will open up a new dimension for the bottom-up synthetic cell assembly. All in all, this shows the great potential of DNA nanotechnology for the development of engineerable and fully-synthetic cells with applications in biomedicine, cellular biophysics and synthetic biology.

## Methods

### DOSU design and assembly

The DNA sequences for the DOSU were adapted from Mohammed et al.^25^. The design is shown in Supplementary Fig. 1. To allow binding of the DOSU to the lipid membrane, we incorporated 12 single-stranded overhangs at columns 1 or 12 and designed complementary cholesterol-tagged DNA linkers (Supplementary Fig. 1a). Moreover, we added 6 biotin-modifications on overhangs at column 6 in order to induce the DOSU clustering with streptavidin (Supplementary Fig. 1b). Typically, 50 to 100 μL of the DOSUs were assembled in 1x tris-acetate-EDTA (TAE) buffer and 12 mM MgCl_2_ by using 16 nM of p7429 scaffold (tilbit), 160 nM of staples and 80 or 160 nM of Atto647N-tagged DNA. The solution was annealed using a thermocycler (BioRad) by heating the solution to 90 °C and subsequently cooling it in steps of 1 °C every minute until holding it for 1 h at a temperature of 45 °C. Subsequently, the solution was further cooled to 32 °C in steps of 0.1 °C. Following the annealing of the DOSU, it was purified three times by spin filtration with an Amicon 50 kDa filter. 50 μL of the annealed solution were diluted with 450 μL of folding buffer (1x TAE, 12 mM MgCl_2_) and centrifuged for 5 min at 10 770 g. This process was repeated three times. After annealing and purification the DOSU concentration was measured with a spectrophotometer and stored at 4 °C for up to three days. The DNA strands were either purchased from Integrated DNA Technologies or Biomers (purification: standard desalting for unmodified DNA oligomers, HPLC for DNA oligomers with modifications). All DNA sequences are listed in Supplementary Table 1 and the Supplementary Data.

### Confocal fluorescence microscopy

A confocal laser scanning microscope LSM 900 (Carl Zeiss AG) was used for confocal microscopy. The pinhole aperture was set to one Airy Unit and the experiments were performed at room temperature. Images of DNA filaments in Figure 4 were acquired using the Airyscan mode. The images were acquired using a 20× (Plan-Apochromat 20×/0.8 Air M27, Carl Zeiss AG) or 63× objective (Plan-Apochromat 63×/1.4 Oil DIC M27). Images were analyzed and processed with ImageJ (NIH, brightness and contrast adjusted).

### Agarose gel electrophoresis

DOSU assembly was verified using agarose gel electrophoresis. 0.7% agarose gel was casted using 0.5x TAE buffer containing 12 mM MgCl_2_. The same buffer was used as running buffer. 10 μL of purified origami (200 ng μL^-1^) was mixed with 6x Blue Loading Dye (New England Biolabs, MA, USA) and loaded into the gel pocket. Quick-Load^®^1 kb Extend DNA Ladder (New England Biolabs, MA, USA) was used as reference. The gel was run on ice at constant voltage of 60 V for 4-5 h and subsequently stained with GelRed^®^(Sigma-Aldrich) then imaged using Azure 600 imager (Azure biosystems).

### Preparation of small unilamellar vesicles

Lipids were stored in chloroform at −20 °C and used without further purification. Small unilamellar vesicles (SUVs) were formed by mixing the chloroform-dissolved lipids (69 %1,2-dioleoyl-sn-glycero-3-phosphocholine (DOPC, Avanti Polar Lipids), 30 %1,2-dioleoyl-sn-glycero-3-phospho-(1’-rac-glycerol) (sodium salt, DOPG, Avanti Polar Lipids) and 1%Atto488/Atto390-1,2-dioleoyl-sn-glycero-3-phosphoethanolamine (Atto488- or Atto390-DOPE, Atto TEC)) or without any fluorophore modification in a glass vial. The lipid solution was dried under a stream of nitrogen gas. To remove traces of solvent, the vial was kept under vacuum in a desiccator for at least 20 min. Lipids were resuspended in 1x DPBS at a final lipid concentration of 2.5 or 5 mM. The solution was vortexed for 10 min to trigger vesicle formation. Subsequently, vesicles were extruded to form homogeneous SUVs with eleven passages through a polycarbonate filter with a pore size of 50 nm (Avanti Polar Lipids, Inc.). SUVs were stored at 4 °C for up to a week or used immediately for GUV formation.

### Cryo electron microscopy

Samples were prepared for cryo-EM by applying 5 μL of sample solution (1x PBS, 10 mM MgCl_2_, SUVs (455 μM lipids) and DOSU) onto a glow-discharged 300 mesh Quantifoil holey carbon-coated R3.5/1 grid (Quantifoil Micro Tools GmbH, Großlöbichau). The grid was blotted for 3 s and plunge-frozen in liquid ethane using a Vitrobot Mark IV (FEI NanoPort, Eindhoven, The Netherlands) at 100 %humidity and stored under liquid nitrogen. Cryo-EM specimen grids were imaged on a FEI Tecnai G2 T20 twin transmission electron microscope (FEI NanoPort, Eindhoven, The Netherlands) operated at 200 kV. Electron micrographs were recorded with an FEI Eagle 4k HS, 200 kV CCD camera with a total dose of ≈40 electrons/Å^2^. Images were acquired at 50000x nominal magnification.

### Image analysis of DOSU insertion angle

The insertion angles to the SUVs of the DOSUs were measured from cryo-EM micrographs using the Angle Tool from ImageJ^29^. The insertion angle is formed by a tangent line to the surface of the SUV and a line parallel to the DOSU. DOSUs are considered to be inserted correctly if the insertion angle is greater than 75° and lower than 105°. The angle histogram of DOSU insertion (Fig. 1**c**) is plotted using Python^30^ (code provided in Supplementary Data). The histogram of DOSU insertion (Supplementary Fig. 6) is plotted using GraphPad Prism.

### Preparation of giant unilamellar vesicles using electroformation

Giant unilamellar vesicles were prepared using the electroformation method^31^ using a VesiclePrepPro device (Nanion Technologies GmbH). 30 μL of 5 mM lipid mix (containing 99 %or 99.5 %1,2-dioleoyl-sn-glycero-3-phosphocholine (DOPC) and 1 %or 0.5 %1,2-dioleoyl-sn-glycero-3-phospho ethanolamine-Atto488 (Atto488-PE) or no fluorophore-modified lipid) in CHCl_3_ were homogeneously spread on the conductive side of an indium tin oxide (ITO) coated glass slide (Visiontek Systems Ltd). After evaporating the chloroform for 20 min under vacuum, a rubber ring was placed on the lipid-coated ITO slide and filled with 270 μL of 270 mM sucrose solution to match the omsolarity of the phosphate buffered saline buffer. The second ITO slide was put on top and the chamber connected to the electrodes of the VesiclePrepPro (Nanion). An AC field (3 V, 5 Hz) was applied *via* the electrodes for 138 min while the solution was heated to 37 °C. GUVs were collected immediately after electroformation and stored at 6 °C for up to 7 days.

### Dye influx assays

GUVs were prepared using electroformation from a lipid mixture containing 99.5 %DOPC and 0.5 %fluorescent Atto488-DOPE lipids. The DOSU was mixed in a ratio of 0.96: 0.02: 0.02 with two types of cholesterol-tagged DNA (100 μM) in order to modify the DOSU with cholesterol. The mixture of DOSU and cholesterol-tagged DNA was incubated for 10 min at 35 °C and 200 rpm. 7.5 μL of electroformed GUVs were diluted in 34 μL 1x DPBS containing 5 mM MgCl_2_ and incubated them with 5 μL cholesterol-tagged DOSU for 1 min to allow binding to the GUVs. Subsequently, 1 μL of 1: 20 diluted AlexaFluor 647 NHS-ester dye (10 μM) was added. At each time point 5-7 μL were pipetted out of the PCR tube and imaged within a custom-built observation chamber using a confocal microscope.

### DNA tile design and filament assembly

DNA filament sequences were adapted from Rothemund et al.^27^. The individual DNA oligomers (5 per tile) were mixed to a final concentration of 5 μM in 1x Dulbecco’s phosphate buffered saline (DPBS) and 10 mM MgCl_2_. 50 to 200 μL of the solution were annealed using a thermocycler (Bio-Rad) by heating the solution to 90 °C and cooling it to 25 °C in steps of 1 to 0.5 °C for 2 or 4.5 h, respectively. The assembled DNA filaments were stored at 4 °C and used within a week after annealing. The DNA strands were either purchased from Integrated DNA Technologies or Biomers (purification: standard desalting for unmodified DNA oligomers, HPLC for DNA oligomers with modifications). All DNA sequences are listed in Supplementary Tables 2, 3.

### Filament disassembly by invader strand influx

DNA signalling units were assembled with adapter tiles and cholesterol-tags and used within one week. Cy3-labeled DNA filaments were assembled at a concentration of 5 μM and designed such that they do not bind to the DOSU. DNA filaments were encapsulated into GUVs at 500 nM in a 1x DPBS inner solution with 10 mM MgCl_2_ using the droplet-stabilized GUV formation method^32^. The iso-osmotic release buffer contained the same MgCl_2_ concentration of 10 mM to stabilize DNA nanostructures on the outside of GUVs. After releasing the DNA filament-containing GUVs, they were reannealed in the thermocycler in steps of 1 °C per 45 s from 90 °C to 25 °C in order to ensure filament assembly within GUVs. Subsequently, 45 μL filament-encapsulating GUVs were mixed with 5 μL cholesterol-tagged DOSU. After incubation for 1 min to allow DOSU binding to GUVs, 2.5 μM invader strand solution was added to disassemble DNA filaments. In the case of the biotinylated clustered DOSU, an additional 2.5 μL streptavidin (5 mg/ml in DPBS was added to the DOSU-GUV mixture after 1 min. For the negative controls either the addition of the DOSU or invader strand was omitted. At each time point 5-7 μL were pipetted out of the PCR tube and imaged within a custom-built observation chamber using a confocal microscope.

### Preparation of giant unilamellar vesicles containing DNA filaments

Giant unilamellar vesicles (GUVs) containing DNA filaments were formed using the droplet-stabilized GUV formation method^32^. Briefly, 1.25 mM SUVs, 5 μM DNA filaments, 10 mM MgCl_2_ and phosphate buffered saline (DPBS consisting of 137mM NaCl, 2.7mM KCl, 10 mM Na_2_HPO_4_ and 1.8 mM KH_2_PO_4_) were mixed together. The aqueous mix was layered on top of an oil-surfactant mix containing 1.4 wt% perfluoropolyether–polyethylene glycol (PFPE–PEG) fluorosurfactants (Ran Biotechnologies) and 10.5 mM PFPE–carboxylic acid (Krytox, MW, 7000–7500 g/mol, DuPont) in a microtube (Eppendorf). The ratio in between aqueous and oil phase was 1:3, generally leading to volumes of 100 μL:300 μL. Droplet-stabilized GUVs were generated by shaking the microtube vigorously by hand. The water-in-oil emulsion droplets were left at room temperature for 0.5 – 1h. Within this incubation period, the SUVs fused at the droplet periphery to create a spherical supported lipid bilayer, termed droplet-stabilized GUV. Afterwards, the oil phase was removed and 100 μL of 1x DPBS was added on top of the emulsion droplets. The droplet-stabilized GUVs were destabilized by addition of 100 μL of perfluoro-1-octanol (PFO, Sigma-Aldrich) to release freestanding GUVs into the PBS. GUVs were stored for up to two days at 6 °C. GUVs were imaged in a custom-built observation chamber that was coated with 2 mg/ml bovine serum albumin (Sigma Aldrich) for 15 min to prevent fusion of the GUVs with the glass coverslide.

### Image analysis of the state of filament assembly

For the filament assembly state analysis, confocal images of the encapsulated filaments were processed with Fiji. First, the Auto threshold was automatically set by the Triangle built-in method and a binary image was generated. The choice of the automatic threshold method triangle was mainly determined by the fact that it supports the maintenance of high intensity connections between adjacent pixels, which is the case for filaments. Second, the built-in Erode function was used to eliminate single non-connected pixels. Depending on the recorded fluorescent intensities this step was applied once or twice resulting in visualizing assembled filaments in contrast to their background or disassembled filaments. Third, the filter performance was checked manually and the image was skipped in case the the filter did not perform well (e.g. intensities of disassembled filaments were too high and close by such that step one and two generate GUVs full of filaments from nothing). If the filter performed well the ROI was set manually matching the GUV size and by the help of a bright field channel. The area fraction of pixels of the processed filament channel was measured within the ROI via the Fiji built-in area fraction measurement option. The results of 15 GUVs or more are averaged for each time step. For more details and processed image examples see Supplementary Fig. 14.

### Filament remodelling experiments

100 μL of GUVs containing 500 nM Cy3-labeled DNA filaments were mixed with 2 μL DNase (4 units, Sigma Aldrich). GUVs were then incubated at 37 °C for 20 min to digest DNA filaments present in the outer aqueous solution due to an imperfect release. The solution was then heated to 95 °C to inactivate the DNase and slowly cooled to 25 °C in steps of 1 °C for 2 h. Subsequently, 2 μL of DOSU was incubated with cholesterol-tagged DNA and added to 36 μL of GUVs. After 1 min of incubation to allow DOSU attachment, 2 μL of streptavidin (Sigma Aldrich, 5 mg/ml) was added. For the negative control the addition of streptavidin was omitted. GUVs were imaged in the equatorial plane after 16 h. The center-of-mass displacement of DNA filaments was analyzed by choosing the GUV center as center (0,0) of a two-dimensional coordinate system. DNA filament fluorescence was thresholded to remove background fluorescence of unpolymerised DNA tiles and the center-of-mass (x,y) of the fluorescence intensity analyzed. The absolute value of the DNA filament center-of-mass was then calculated in the reference of the GUV according to 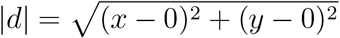.

### Statistical analysis

All the experimental data were reported as mean ±SD from n experiments, filaments or GUVs. The respective value for n is stated in the corresponding figure captions. All experiments were repeated at least twice. In Fig. 1**c** the number of analysed individual SUVs is *n* = 287. In Fig. 1**f** the number of analysed individual GUVs per timestep is in the range of 20 < *n* < 24 (*n* = 150 in total) for GUVs with DOSU (upper plot) and 30 < *n* < 32 (*n* = 216 in total) for plain GUVs (lower plot). In Fig.2**d** the number of analysed individual GUVs per time step is in the range of 39 < *n* < 44 (*n* = 567 in total). In Fig. 3**c** the number of analysed individual GUVs for each time point is *n* = 20 (*n* = 140 in total). In Fig. 3**e** the number of analysed individual GUVs is as follows: - 0 h *n* = 34,-5 h *n* = 28, + 0 h *n* = 31, +5 h *n* = 69. In Fig. 4**c** the numbers of analyzed DNA filaments are *n* = 628 and *n* = 162 and individual tiles (“ssDNA”) are *n* = 186 and *n* = 196, respectively. In Fig. 4**e** the number of analysed individual GUVs is *n* = 10 per condition (*n* = 20 in total). To analyze the significance of the data, a Mann-Whitney test was performed using Prism GraphPad (Version 9.1.2) and p-values correspond to ****: p ≤ 0.0001, ***: p ≤ 0.001, **: p ≤ 0.01, *: p ≤ 0.05 and ns: p ≥ 0.05.

## Funding

K.G. received funding from the Deutsche Forschungsgemeinschaft (DFG, German Research Foundation) under Germany’s Excellence Strategy via the Excellence Cluster 3D Matter Made to Order (EXC-2082/1 - 390761711), the Hector Fellow Academy and the Max Planck Society. K.J. thanks the Carl Zeiss and the Joachim Herz Foundation for financial support. The Max Planck Society is acknowledged for its general support.

## Author contributions

K.J. and K.G. designed the study. K.J. developed transmembrane DOSUs and proved dye and ssDNA influx into GUVs. M.I. performed and analyzed dye and single-stranded DNA influx experiments. M.S. conducted DOSU characterization experiments. K.J. performed filament disassembly and displacement experiments. M.I. analyzed filament disassembly experiments. K.J. analyzed filament displacement experiments. K.J. designed and verified clustered DNA DOSU assembly. K.J. and M.I. proved the enhanced DNA DOSU insertion efficiency for clustered DOSU. K.J. supervised M.I. and M.S. during the course of this study. U.M. performed Cryo-EM imaging. M.T. analyzed cryo-EM images. K.J., M.I. and K.G. wrote the manuscript with the help from all authors.

## Competing interests

The authors declare no competing interests.

## Data availability

The datasets generated during and analysed during the current study are available from the corresponding author on reasonable request.

## Additional information

Correspondence and requests for materials should be addressed to K.G.

## Notes

### Competing Interest Statement

The authors have declared no competing interest.

