## Supplementary Information for "Mechanochemical signal transduction in synthetic cells"

*

**References** **24**

### Supplementary Notes

#### Approximation of DOSU density on GUVs

In order to quantify the density of the DOSUs on GUVs, the number of GUVs per 1 µL sample were estimated from fluorescent confocal images (compare Supplementary Fig. 7**a,b,c**). For this, we assumed even partitioning of the DNA origami on the available membrane surface area.Very small GUVs with a cross sectional area in the equatorial plane below 30 µm$^2$ were not counted. From the confocal plane with the maximal cross sectional area of a GUV $A_{\mathrm{cs}}$ the radii $r = \sqrt{A_{\mathrm{cs}}/\pi}$ and surface areas $A_{\mathrm{s}} = 4 \cdot A_{\mathrm{cs}}$ were calculated, respectively (with an estimated uncertainty of 1 %). The absolute numbers obtained from two images, SupplementaryFig. 7**a,b** were averaged. The height of the observation chamber from both images were evaluated manually from z-projections and averaged. Hereby we obtained a total volume of $(638.9 \times 638.9 \times 97)$µm$^3$ and therefore calculated the GUV density to be $2898 \pm 205$ GUVs/µL. The resulting average surface area $A_{\mathrm{s}}$ amounts to $(74900 \pm 749)$ µm$^2$.

The DNA origami (Supplementary Fig.7) was diluted 1:100 resulting in a dsDNA concentration of 2.27 ng/µL as measured with a photospectrometer (IMPLEN, NanoPhotometer NP80). With a particle analysis tool (Analyze Particles from ImageJ), we measured 1938, 1885, 1812 particles for lower particles size thresholds 5, 10, 15 pixels, respectively, in a sample volume of $(77.34 \times 77.34 \times 97)$µm$^3$ ($V$ =0.000 58 µL). Averaging these three values leads to a density of $(32.3 \pm 1.1)10^5$ particles/µL for the undiluted, purified DOSU. For the experiments the DOSU was mixed with cholesterol-tagged DNA and diluted such that a final concentration of (21.8, 4.4, 0.9) ng/µL (high, middle, low) of the DOSU solution was present in the final sample. Including this into the approximation, we obtain the following density of DOSUs per GUV: $(16.37 \pm 1.00, 3.28 \pm 0.20, 0.65 \pm 0.04)$ DOSUs/µm$^2$, respectively. In absolute numbers, this corresponds to approximately $10700 \pm 1200$, $2100 \pm 200$ and $400 \pm 50$ DOSUs per GUV, respectively. Their average distance on the GUV membrane is 265 nm, 590 nm, 1 µm (high, middle, low, respectively).

### Supplementary Figures

#### Supplementary Figure 1: Functionalized DOSU design

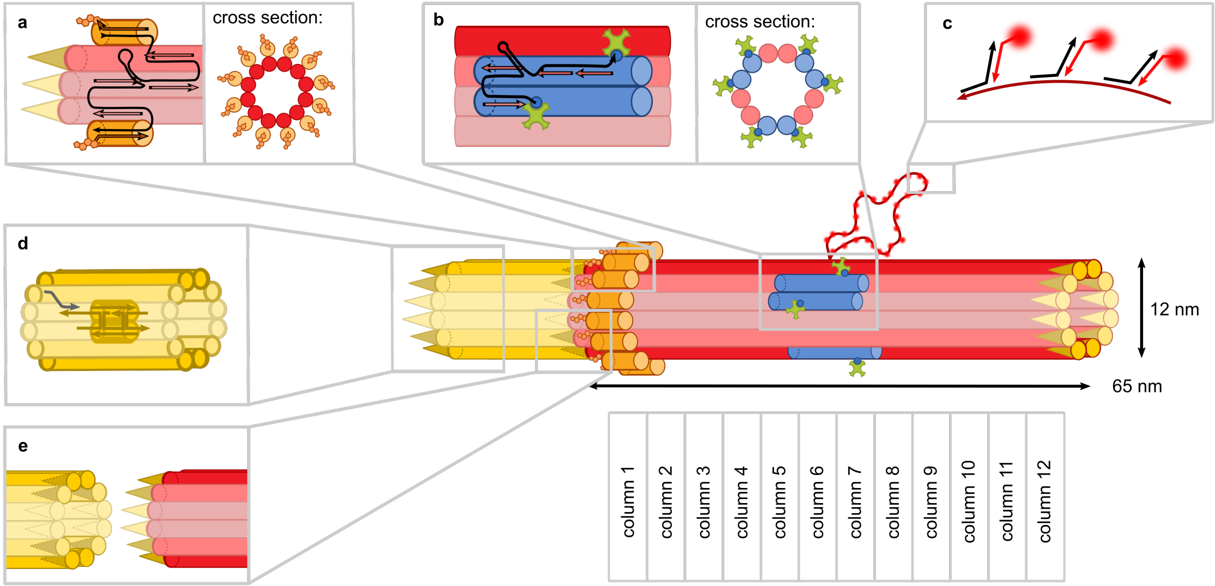

Figure 1: Sketch of the modified and functionalized DOSU design (basic design adapted from Mohammed et al.[1]). The DOSU consists of 12 parallel double helices that are sketched as rods and form a hollow cylindrical DNA origami structure. Numerically, the DNA origami measures 12 nm in outer diameter and 65 nm in length. The design features 12 repetitive columns containing 6 staple strands each that hybridize with the scaffold strand (termed column 1-12). **a** The replacement of the six staple strands in column 1 by modified strands with overhangs on their 5' and 3' ends allows 5'- and 3'-cholesterol-functionalized DNA strands to hybridize to the DOSU. This modification results in 12 bound hydrophobic cholesterols in column 1 evenly distributed around the circumference of the DNA origami as sketched in the cross sectional view. Modifying the DOSU with hydrophobic moieties enables its insertion into the lipid membrane. **b** Three of the six staple strands on column 6 were modified with biotin on their respective 3' as well as the 5' end. This adds up to six biotins distributed over column 6 and 7 around the DOSU as sketched in the cross sectional view and enables programmable multimerization of the DNA origami in the lateral dimension. **c** An overhanging scaffold loop located at the center of the DOSU hybridizes with 100 overhang staple strands, which bind complementary fluorescently-labeled strands (Atto647-labeled, $\lambda_{ex}$ =647 nm) enabling visualization of single DOSUs with confocal microscopy.[1]

Figure 1: **d** DNA tiles can be designed to form DNA nanotubes. This original DNA filament tile system can be modified with a sticky end in order to allow an invader strand to disassemble the filaments into single non-binding tiles. **e** The DOSU can be functionalized with a DNA-based adapter to bind to the DNA filaments.[1] In this way, DNA filaments can grow from the DOSU.

#### Supplementary Figure 2: Bulk confocal images of DOSUs

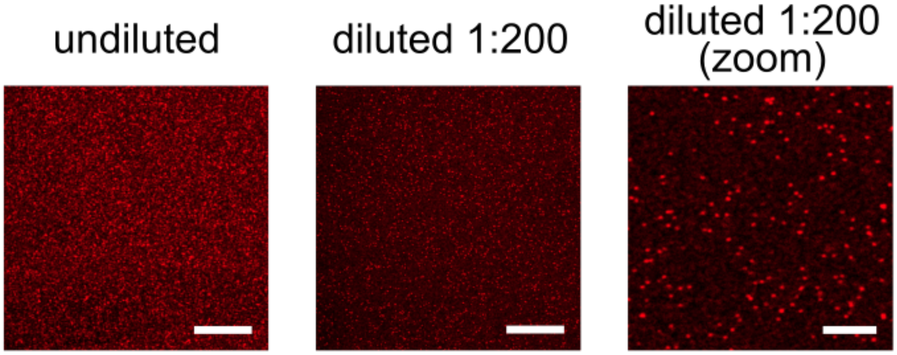

Figure 2: Bulk confocal images of DOSUs (after purification by spin filtration). Confocal images of undiluted and diluted (1 to 200) DOSU (Atto647-labeled, $\lambda_{ex}$ =647 nm) in bulk bound to glass coverslides. The DNA concentration after purification was measured to be 180 ng/µL by UV-vis spectrometry. Because of the high number of fluorophores (100 per DNA origami), individual DOSUs can be visualized with confocal microcopy indicating the successful folding. Scale bar: 10, 20 and 5 µm, respectively.

#### Supplementary Figure 3: Self-assembly of DOSUs

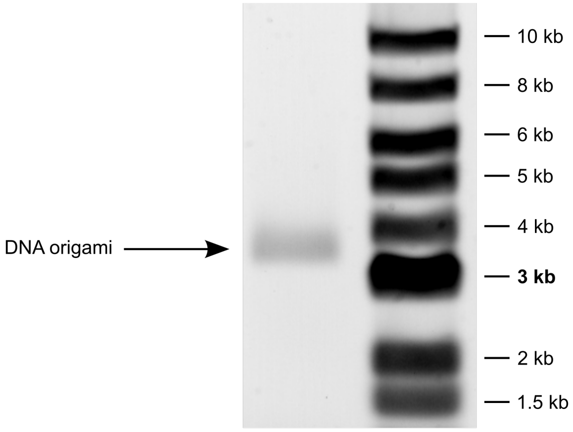

Figure 3: Agarose gel electrophoresis of the DOSUs after purification. The DNA origami band is shown on the left lane and the DNA ladder is shown on the right lane. The molecular weight of the ladder is indicated in kilo base pair (kbp).

#### Supplementary Figure 4: Stability of DOSUs

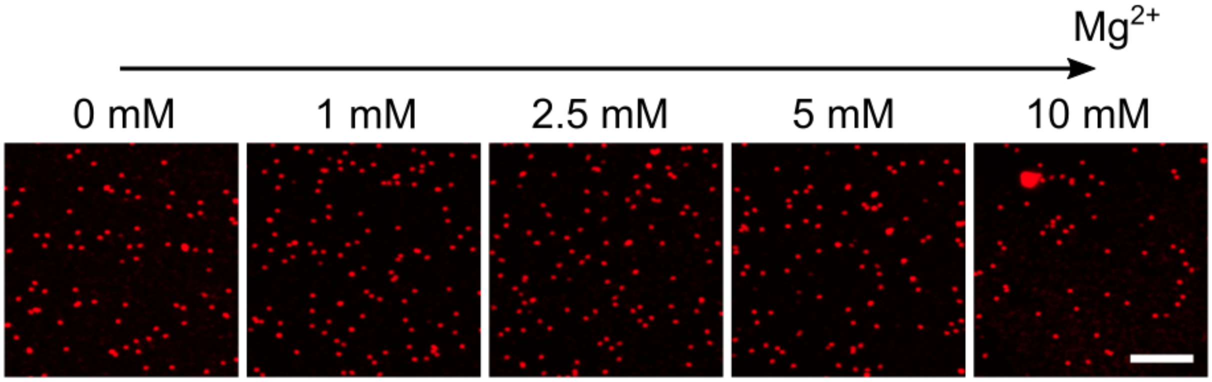

Figure 4: Bulk images of purified DOSUs at different concentrations of magnesium ions 72 h after purification. Confocal images of DOSUs (Atto647-labeled, $\lambda_{ex}$ =647 nm) in bulk bound to glass coverslides. DOSUs in 1x PBS are stable after three days even in the absence of magnesium ions. Scale bar: 5 µm.

#### Supplementary Figure 5: Cryo electron micrographs of DOSUs inserting into the membrane of SUVs

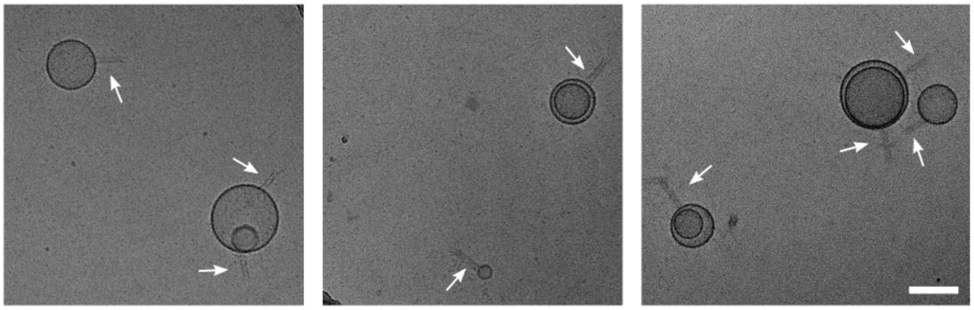

Figure 5: Cryo electron micrographs of DOSUs inserting into the membrane of SUVs (DOSUs are highlighted with white arrows). Scale bar: 100 nm.

#### Supplementary Figure 6: Insertion of DOSU on SUV

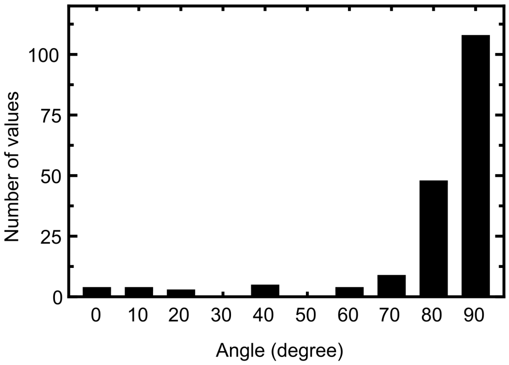

Figure 6: Histogram with the distribution of insertion angles of DOSUs on SUV. The insertion angles are analyzed from cryo-EM micrographs (as described in the Methods section). An angle histogram is also plotted from the same data in Figure 1.

#### Supplementary Figure 7: DOSU density on GUV membrane

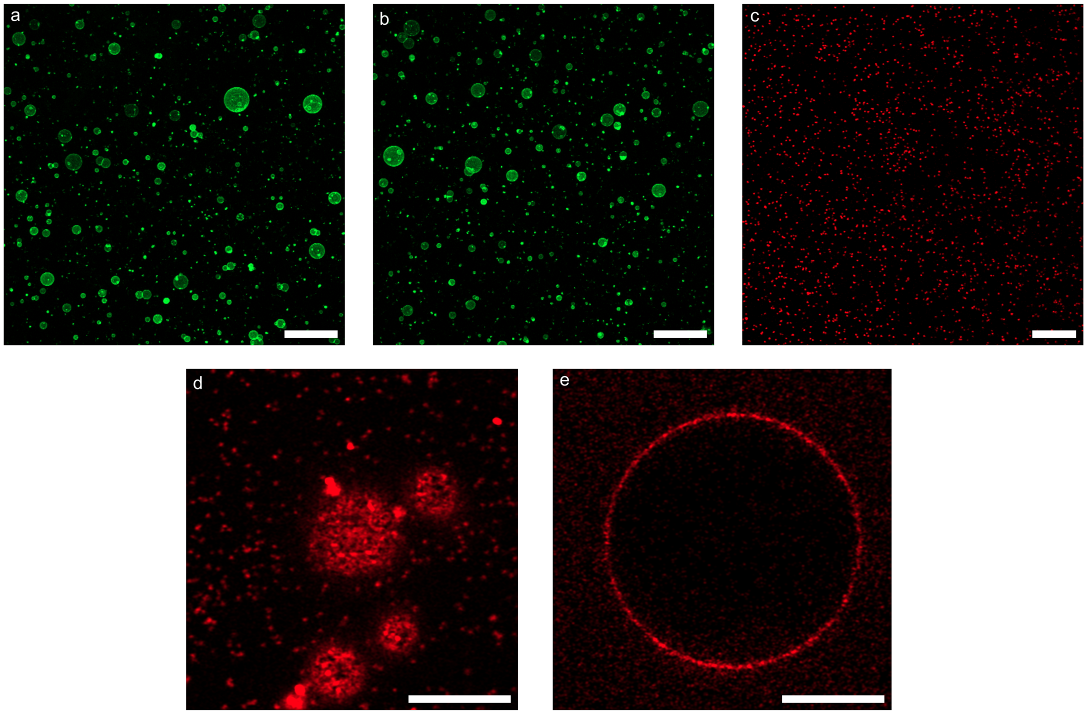

Figure 7: To estimate the DOSU density on GUV membranes fluorescent confocal images of GUVs and DOSUs have been analyzed. **a** + **b** Analysed maximum intensity z projections of GUVs. Scale bar: 100 µm. **c** Confocal image of DOSUs (Atto647-labeled, $\lambda_{ex}$ =647 nm). Scale bar: 10 µm. **d** + **e** Confocal images of DOSUs (Atto647-labeled, $\lambda_{ex}$ =647 nm) on the GUV membrane (unlabeled) of four single GUVs at the lowest z-position (**d**) and at the central slice of one single GUV (**e**). From this data, the DOSU densities on the GUV membrane have been estimated as described in the Supplementary Note.

#### Supplementary Figure 8: DOSU transmembrane insertion is controlled by cholesterol-tagged DNA

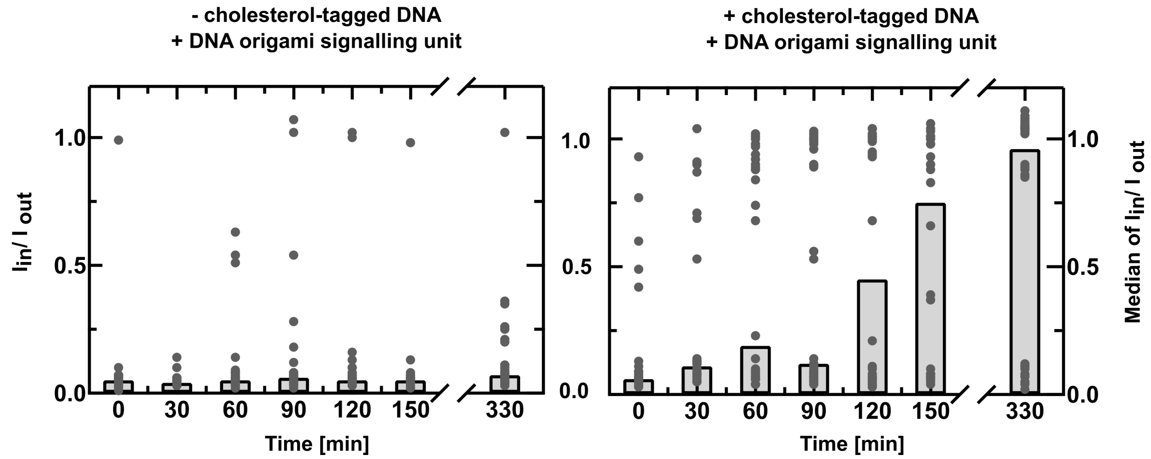

Figure 8: DOSU transmembrane insertion is controlled by cholesterol-tagged DNA. Ratio of inner intensity $I_{\text{in}}$ over outer intensity $I_{\text{out}}$ of individual GUVs and their median value over time for GUVs with high concentration of DOSUs (21.8 ng/µL) and in absence of cholesterol tagged DNA (left, in total $n = 208$) and in presence of cholesterol-tagged DNA (right, in total $n = 210$). The experiment indicates that the modification of the DOSU with cholesterol tags (Supplementary Fig. 1**a**) leads to transmembrane insertion and thereby allows for influx of the fluorescent dye into the GUV lumen.

#### Supplementary Figure 9: DOSU-mediated dye influx into GUVs

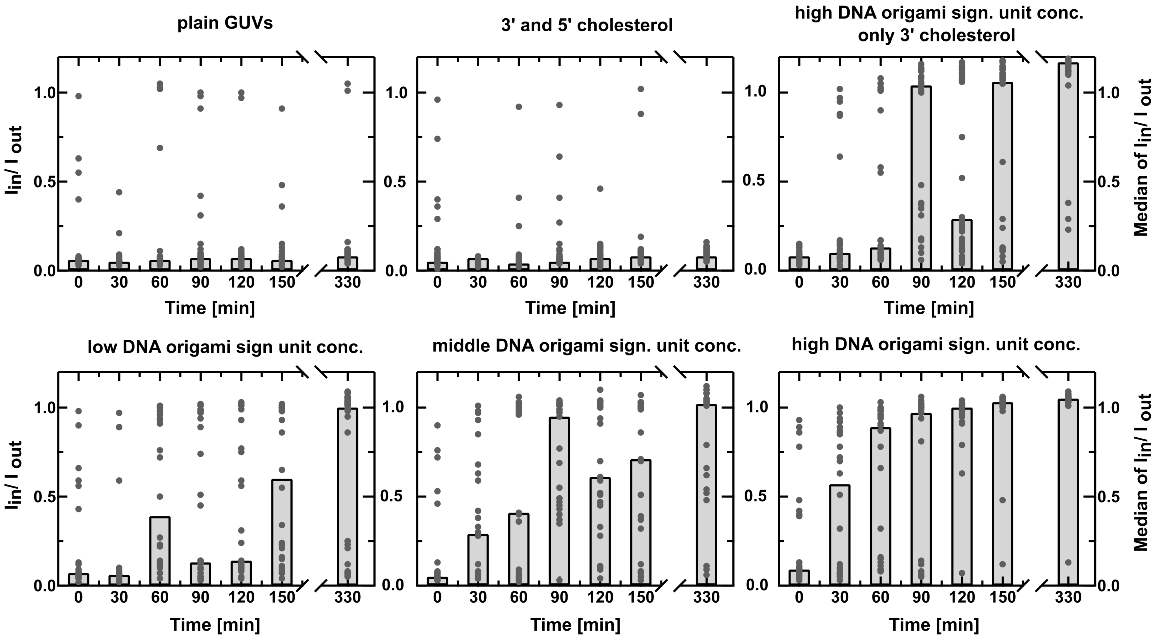

Figure 9: DOSU-mediated dye influx into GUVs. Ratio of inner intensity $I_{\text{in}}$ over outer intensity $I_{\text{out}}$ of single GUVs ($n = 30$) and their median value over time. Different DOSU concentrations are compared to two negative controls (plain GUVs and GUVs with 3'- and 5'-cholesterol-tagged DNA strands). A sample of 50 µL contained 7.5 µL GUVs, 10 µM fluorescent dye (Atto647-labeled, $\lambda_{ex}$ =647 nm). 5 µL, 1 µL, 0.25 µL DOSU-cholesterol solution respectively (dsDNA density high: 21.8 ng/µL, middle: 4.4 ng/µL, low: 0.9 ng/µL) in a 1x PBS buffer with 5 mM $MgCl_2$. One sample contained only half of the cholesterol tags because only the 3' DNA staple strands were added to the DOSU (see Supplementary Fig. 1 a). The DOSU concentration after purification was 227 ng/µL. Both negative controls without DOSU (plain or with cholesterol incubated GUVs) do not show dye influx over time. This proves that neither the used dye nor cholesterol-tagged single stranded DNA perturbs the GUV membrane permeability. The third plot confirms that half the number of cholesterol tags deteriorates the insertion efficiency compared to plot 6. Moreover, a higher amount of DOSUs also leads to more dye influx as it is shown by the plots in the bottom row. Note that the same amount of osmolarity-matched buffers have been added to all samples to exclude leakage induced by the additions or by osmotic shocks.

#### Supplementary Figure 10: DOSU-mediated dye influx into GUVs over longer periods

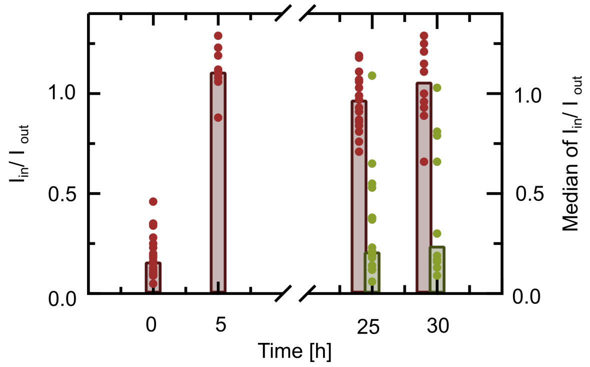

Figure 10: Ratio of inner intensity $I_{\text{in}}$ over outer intensity $I_{\text{out}}$ of single GUVs ($n \geq 9$) and their median value over time is shown. A sample of 50 µL contained 7.5 µL fluorescently labeled GUVs (orange, LissRhod-labeled, $\lambda_{ex}$ =561 nm), 5 µL DOSU with cholesterol tags and 10 µM fluorescent dye (red, Atto647-NHS ester, $\lambda_{ex}$ =647 nm) in 5 mM $MgCl_2$ and 1x PBS buffer. After 24 h 10 µM second fluorescent dye (green, Alexa Fluor 488 Carboxylic acid, $\lambda_{ex}$ =500 nm) was added and single GUVs were analyzed on both fluorescent channels (in total 4 measurement were performed). Images with very small GUVs ($< 5um$) excluded from the analysis and all images were thresholded for $I_{in}/I_{out} < 1.3$. For better visualization, both, the 24 h and 29 h dataset was separated into a first (red) and second (green) dataset and plotted next to each other. We observe several single GUVs that show influx of both dyes and conclude that the DOSU-mediated membrane permeation is still maintained, suggesting a non-transient insertion.

#### Supplementary Figure 11: DOSU-mediated dye efflux from GUVs

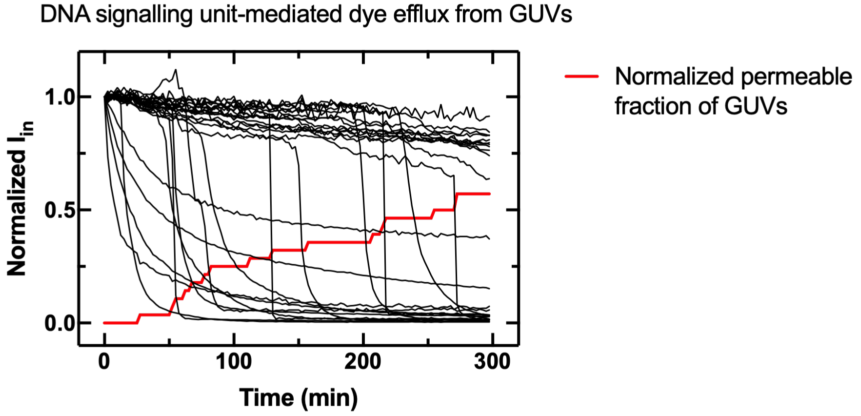

Figure 11: DOSU-mediated dye efflux from GUVs. Normalized inner intensity of GUVs (black) containing 50 µM membrane-impermeable dye (Atto488-NHS ester, $\lambda_{ex}$ =488 nm) with attached DOSUs (Atto647-labeled, $\lambda_{ex}$ =647 nm) incubated with cholesterol-tagged DNA obtained from confocal images. After insertion of DOSUs the dye flows out of the GUV within a few minutes. The normalized permeable fraction of GUVs (red) increases over time and represents the fraction of GUVs with $I_{\text{in}} < 0.2$.

#### Supplementary Figure 12: DOSU clustering via streptavidin

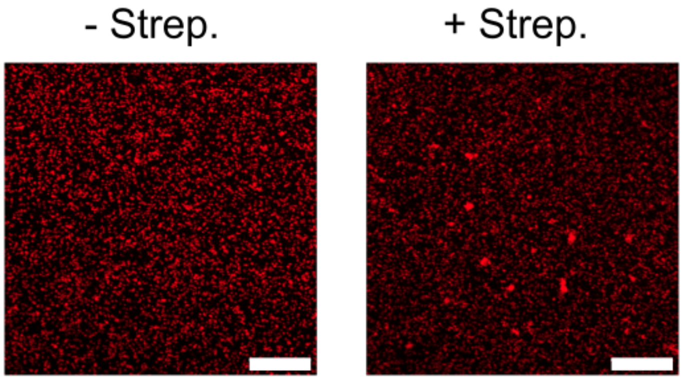

Figure 12: DOSU clustering via streptavidin. Confocal images of DOSUs (Atto647-labeled, $\lambda_{ex}$ =647 nm) incubated with cholesterol-tagged DNA in presence and absence of 250 µg mL$^{-1}$ in 1x PBS and 5 mM $MgCl_2$ after 330 min of incubation. Scale bar: 10 µm.

#### Supplementary Figure 13: DOSU-mediated single stranded DNA influx into GUVs

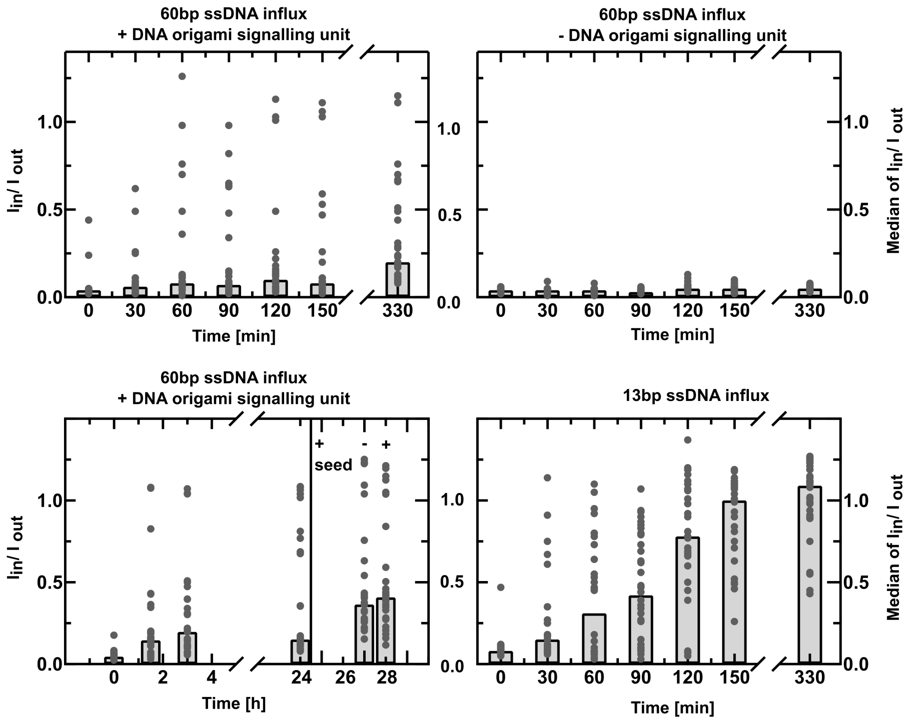

Figure 13: DOSU-mediated single stranded DNA (ssDNA) influx into GUVs. A sample contained unclustered DOSUs, GUVs and either 62 base pairs (bp) long ssDNA (Cy3-labeled, $\lambda_{ex}$ =561 nm) shown in the first three graphs or 13 bp long ssDNA (6T-2A-5T Atto647-NHS ester-labeled, $\lambda_{ex}$ =647 nm) shown in the forth graph, as indicated. The first two plots depict the ratio of inner intensity $I_{\text{in}}$ over outer intensity $I_{\text{out}}$ of single GUVs ($n = 30$) as well as a histogram of their median values with (left) and without (right) DOSU, respectively. The third plot shows a long time study of the 62 bpssDNA influx into GUVs with a second addition of DOSU after 24 h of incubation an insignificant change of the distribution of $I_{\text{in}}/I_{\text{out}}$ of the analyzed single GUVs ($n = 30$). The fourth plot shows a shorter ssDNA influx study for the same DOSU over time ($n = 30$), which has an increased influx into GUVs due to the smaller size of the ssDNA.

#### Supplementary Figure 14: Analysis of clustered DOSU-mediated chemical signalling to disassemble DNA cytoskeletons

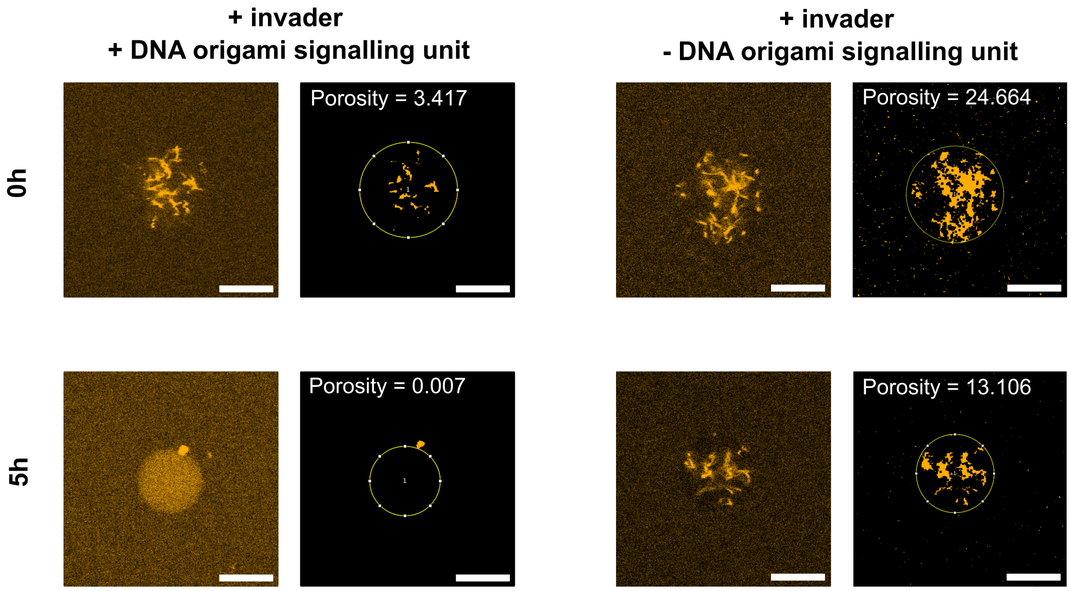

Figure 14: Analysis of clustered DOSU-mediated chemical signalling to disassemble DNA cytoskeletons. Four confocal images of single GUVs with DNA cytoskeletons (Cy3-labeled, $\lambda_{ex}$ =561 nm) and their processed version are shown in presence of invader strand (5 µM at the outside and in presence (left) and absence (right) of DOSU for 0 h and 5 h. Scale bar: 10 µm. The processed image shows the thresholded images to analyse the assembly state of the DNA cytoskeletons. After thresholding and processing the measured porosity value of the selected region of interest depicts whether the filaments are assembled (high value 1) or disassembled (low value ≈ 0). The analysis script is adapted with one (left column), two (right column) or three repeats of applying the erode function to eliminate single artefacts and yield connected pixel accumulations (filaments) regarding different intensities of the sample while the confocal microscopy settings have been fixed beforehand. The choice of the script's adaptation version is clearly determined by the optimum of a complete dataset which includes two timepoints and two measurements of $n$ = 20 single GUVs each. The porosity values were normalized to the initial measurement of the negative control. Regions of interest are selected manually and images are skipped in case the filter did not perform well because of any artefacts. In presence of the engineered clustered DOSU we observe the disassembly of cytoskeletons (++, 5 h, porosity=0.007) and thereby chemical signal transduction across the GUV membrane.

#### Supplementary Figure 15: DOSU-mediated chemical signalling to disassemble DNA cytoskeletons

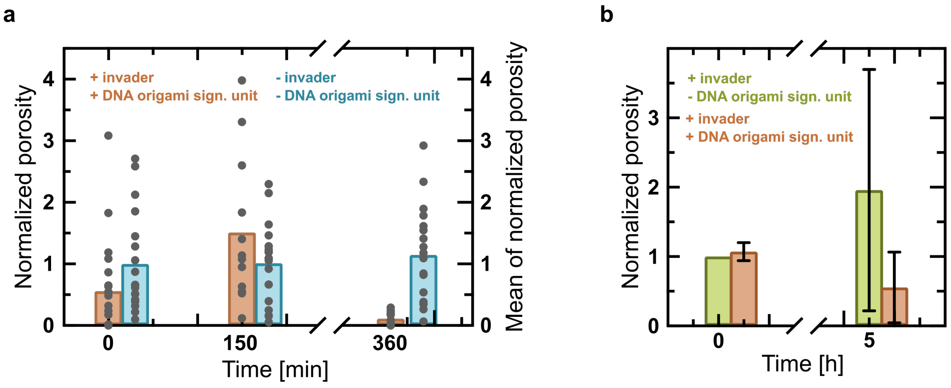

Figure 15: DOSU-mediated invader strand influx into GUVs to disassemble DNA cytoskeletons at the inside of GUVs (compare Supplementary Fig. 1 **d**). The two plots show the decrease of the porosity of DNA cytoskeletons due to filament disassembly resulting from the DOSU-mediated invader strand influx. We performed the disassembly experiment with the unclustered DOSU. Encapsulated DNA filaments (Cy3-labeled, $\lambda_{ex}$ =561 nm) were imaged with fluorescent confocal microscopy and processed and analysed as described in the Methods section. The porosity of the DNA cytoskeleton describes the state of assembly of DNA filaments, assembled filaments result in a high porosity whereas disassembled filaments (homogeneously distributed single tiles) result in a low porosity. **a** Porosity of DNA cytoskeletons within single GUVs ($n = 20$) after 0 min, 150 min and 360 min in presence of invader strands and DOSU (brown) and in absence of both (blue), normalized to the initial measurement of the latter condition. We observed that DNA cytoskeletons remain stable over time within GUVs without any further perturbation (- invader, - DOSU). In addition we find a significant decrease in normalized porosity in presence of the DOSU at $t = 360$ min. **b** Mean and standard deviation of three technical repeats of the porosity decrease of DNA cytoskeletons which is normalized to the initial measurement in presence of invader strand and absence of DOSU ($n = 20$). Also here we observe a porosity decrease after 5 h in presence of the DOSU and thereby chemical signalling across DOSUs.

### Supplementary Tables

#### Supplementary Table 1: DNA sequences for DOSU clustering

Table 1: DNA sequences from 5' to 3' for DOSU clustering. Note that all DOSU sequences can be found in the Supplementary Data.

| Name | DNA sequence |
|---|---|
| T_1R12E_CYC_HP_biotin | Biotin-TTTTATCACCGGCGAGAGGCTGCGTCGTT-TTCGACGCAGTTCTTTTGCAATCCTGAATTT-Biotin |
| T_1R4E_HP_biotin | Biotin-TTTCAATGACAGCTTGATATGGCGAGCTT-TTGCTCGCCATTCCGATAGTCTCCCTCATTT-Biotin |
| T_1R8E_HP_biotin | Biotin-TTTCCAGGCGCGAGGACAGCTCTGGACTT-TTGTCCAGAGTTATGAACGGGTAGAAAATTT-Biotin |

#### Supplementary Table 2: DNA sequences for single-tile DNA filaments that can undergo toehold-mediated strand displacement

Table 2: DNA sequences from 5' to 3' for st DNA filaments, adapted from.[2]

| Name | DNA sequence |
|---|---|
| st1_1 | CTCAGTGGACAGCCGTTCTGGAGCGTTGGACGAAACT |
| st1_2 | TGGTATTGTCTGGTAGAGCACCACTGAGAGGTA |
| st1_3 | CCAGAACGGCTGTGGCTAAACAGTAACCGAAGCA-CCAACGCTGGTAAGTCTCCTTCTTATCT(-Cy3) |
| st1_4 | CAGACAGTTTCGTGGTCATCGTACCT |
| st1_5 | CGATGACCTGCTTCGGTTACTGTTTAGCCTGCTCTAC |
| invader | ACCAGACAATACCAATCCGC |

#### Supplementary Table 3: DNA sequences for single-tile DNA filaments that bind to the DOSU

Table 3: DNA sequences from 5' to 3' for st DNA filaments, adapted from.[2]

| Name | DNA sequence |
|---|---|
| st2_1 | CGTATTGGACATTTCCGTAGACCGACTGGACATCTTC |
| st2_2 | CTGGTCCTTCACACCAATACGGCATT |
| st2_3 | (Atto488-)TCTACGGAAATGTGGCAGAATCAATCA-TAAGACACCAGTCGG |
| st2_4 | ACCAGGAAGATGTGGTAGTGGAATGC |
| st2_5 | CCACTACCTGTCTTATGATTGATTCTGCCTGTGAAGG |

### Supplementary Videos

#### Supplementary Video 1: DOSU-mediated dye efflux from GUVs

DOSU-mediated dye efflux from GUVs. Confocal time lapse of DOSUs (Atto647-labeled, $\lambda_{ex}$ =647 nm) incubated with cholesterol-tagged DNA on GUVs containing 50 µM membrane-impermeable dye (Atto488-NHS ester, $\lambda_{ex}$ =488 nm). The dye flows out of the GUV within 10 min of the observation. Scale bar: 10 µm.

#### Supplementary Video 2: DNA filament disassembly by invader strand in bulk

DNA filament disassembly by invader strand addition in bulk. Confocal time lapse of DNA filaments (orange, Cy3-labeled, $\lambda_{ex}$ =561 nm) upon addition of invader single strand (10 µM) from the right hand side. The DNA filaments were disassembled within few minutes over a lengths of 100 µm. Scale bar: 30 µm.

#### Supplementary Video 3: Chemo-mechanical signalling cascade across the GUV membrane induces symmetry breaking of DNA cytoskeletons

Chemo-mechanical signalling cascade across the GUV membrane induces symmetry breaking of DNA cytoskeletons. Color-coded confocal z-projection of DNA filaments (Cy3-labeled, $\lambda_{ex}$ =561 nm) bound to transmembrane DOSUs inside a GUV. Scale bar: 10 µm.
